# Sedentary behavior, physical inactivity and body composition in relation to idiopathic infertility among men and women

**DOI:** 10.1101/511030

**Authors:** Foucaut Aude-Marie, Faure Céline, Julia Chantal, Czernichow Sébastien, Levy Rachel, Dupont Charlotte, the ALIFERT collaborative group

**Author notes:** Corresponding author, (AMF). **Preferred citation:** Foucaut A-M, Faure C, Julia C, Czernichow S, Levy R, Dupont C, and the ALIFERT collaborative group. Sedentary behavior, physical inactivity and body composition in relation to idiopathic infertility among men and women.

## Abstract

**Background:** Physical activity and sedentary behavior has inconsistent effects on fertility. High body mass index is associated with infertility but to our knowledge, very few studies have explored body composition in association to fertility.

**Objective:** To assess the association between physical inactivity, sedentary behavior, body composition and idiopathic infertility in French men and women.

**Design:** We conducted a case-control multicentric observational study. 159 infertile (79 men and 80 women) and 143 fertile (72 men and 71 women) were recorded in four fertility centers.

**Main Outcome Measures:** Participants completed self-administered questionnaires on sociodemographic and lifestyle characteristics, dietary intake, physical activity and sedentary behavior. Anthropometrics were measured, and bioelectrical impedance analysis was used to estimate body composition. Multivariable logistic regression was used to analyze the association of fertility with PA level and sedentary behavior.

**Results:** In men, being physically inactive (Odd ratio [OR] 2.20; 95% confidence interval [CI], 1.06, 4.58) and having fat mass greater than the reference values for their age (OR 2.83; 95%CI, 1.31, 6.10) were positively associated with infertility. Sedentary behavior and fat-free mass were not related to infertility in men. In women, sedentary behavior (OR 3.61; 95%CI, 1.58, 8.24), high body fat (OR 3.16; 95%CI, 1.36, 7.37) and low fat-free mass (OR 2.65; 95%CI, 1.10, 6.37) were associated with infertility. PA level was not associated with fertility in women.

**Conclusions:** This study suggests that sedentary behavior and physical inactivity would represent two independent risk factors associated with fertility. The various elements that make up physical activity (frequency, intensity, time, and type of exercise) and the interrupting time spent sitting should be considered. Body composition variation should be explored further in relation to the biological pathways involved in idiopathic infertility. Moreover, the improvement of lifestyle factors should be considered in infertility treatment.

## Introduction

Sedentary behavior and physical inactivity represent major health concerns. Sedentary behaviors are defined as any waking activities characterized by energy expenditure below 1.5 metabolic equivalent task (MET) of sitting or lying down. Physical inactivity represents an insufficient volume of physical activity (PA) in daily life, being a level not reaching the recommended PA (150 minutes of moderate PA per week) [1]. These two behaviors are in some cases coexistent, and sometimes not. An individual may have both sedentary behaviors and be physically active [2,3]. In this case, PA can moderate but not offset the deleterious effects of sedentary behavior [4]. It has been shown that sedentary behaviors and physical inactivity independently influence several health factors, non-communicable diseases and mortality [4–6].

Notably, PA has an inconsistent effect on fertility. In men, moderate PA has been positively associated with semen quality [7–10]. However, it was not associated with higher reproductive success in the context of fertility treatment [10]. Some previous studies failed to demonstrate a relationship between PA and semen quality [11,12]. In women, moderate PA increased fecundity parameters and live birth rates, regardless of body mass index (BMI) [13,14]—even during assisted reproductive treatment [15–17]. However, vigorous activity has been associated with lower semen quality in men [18–20] and decreased fertility in women [21–23]. Notably, sedentary behavior has not been clearly associated with semen quality [10,17,18,24,25], though reduced sperm concentration has been linked to increased television watching [9]. In women, sedentary behavior has not been associated with lower fertility in recent studies [17,26].

Obesity is associated with both sedentary behavior and physical inactivity [27,28]. Being overweight and obese is known to impact the fertility of couples. Large cohort studies showed that a BMI over 25 kg/m² (as estimated by the height/weight^2^ ratio) was linked to infertility in both males and females [29,30]. Obesity has been associated with reduced semen quality [31], sperm concentration [29,32–34], mobility [35], DNA damage [36–38], poor oocyte quality, and impaired ovulation and implantation [30]. In the aforementioned studies, obesity estimation was based on BMI values. However, anthropometrics are not the most sensitive parameters for estimating body composition alterations and linking those to clinical outcomes [39]. To our knowledge, very few studies have explored body composition or adiposity in association with fertility, especially fat mass and fat-free mass parameters. Recent studies have used waist circumference and BMI as proxy measures of body composition [40,41]; one used dual-energy X-ray absorptiometry for fat, fat-free mass, and bone mass in 41 young infertile women [42].

The primary objective of this study was to determine if physical inactivity, sedentary behavior and body composition were related to idiopathic infertility in men and women in a French case-control study of nutritional determinants of idiopathic infertility.

## Materials and Methods

Participants were recruited in the ALIFERT case-control multicentric observational study (“ALImentation et FERtilité”, ClinicalTrials.gov identifier: NCT01093378), which evaluated the associations between nutritional parameters and fertility among infertile and fertile couples. The institutional review board approved the study (ALIFERT study - national biomedical research P071224/AOM 08180: NEudra CT 2009-A00256-51).

Data were recorded from 302 French participants, with included 159 infertile (79 men and 80 women) and 143 fertile (72 men and 71 women). Men under 45 years of age and women under 38 years of age were included. Infertile participants had a history of primary idiopathic infertility for at least 12 months of unprotected sexual intercourse, with no diagnosed etiology for their infertility. Men were excluded if they had severe oligozoospermia (< 5 million/mL), azoospermia, or any abnormality of the male genital tract. Women were excluded if they presented anovulation, ovarian failure, or uterotubal pathology. Fertile participants had a recent natural and spontaneous pregnancy and delivery (< 24 months) with a time to conceive shorter than 12 months. No specific matching was conducted between cases and controls, and one fertile control couple was selected for each case couple.

### Data collection

Participants completed self-administered questionnaires on sociodemographic and lifestyle characteristics (sex, age, educational level, and smoking status), dietary intake (semi-quantitative validated food frequency questionnaire), physical activity and sedentary behavior. Anthropometrics, body composition, and blood pressure were measured using standardized procedures (tensiometer; Omron M5-I). Blood samples were used to evaluate plasma high-density lipoprotein (HDL), triglycerides and fasting glycaemia in mmol/L. Assessments were performed after an 8-hour fasting period.

### Physical activity and sedentary behavior assessment

PA level and sedentary behavior were estimated by the self-administered validated last-7-day International Physical Activity Questionnaire (IPAQ) [43]. PA levels correspond to the PA level of a typical week during the inclusion period. Total PA level scores were expressed in MET per minute per week (MET-min/week), which is a product the intensity, duration and frequency of PA retrieved from the items for moderate PA, vigorous PA (occupational and leisure time) and from walking activities (in min/week). Accordance with guideline targets (150 min/week of moderate-to-vigorous PA) were estimated by adding times of moderate, vigorous and walking activities (in min/week). Sedentary behavior was assessed through a question regarding time spent sitting during typical week days (in h/day). A threshold of 5h per day was chosen to categorize participants as having sedentary behavior (≥ 5h/day) or not (< 5h/day). This threshold corresponds to the average time spent while sitting (when occupational time is included) in the general French population [44].

### Anthropometric and body composition assessment

The height and weight of participants was measured to the nearest 0.5 cm and 0.5 kg, respectively, with participants wearing light clothing and no shoes using standardized procedures. BMI (kg/m²) was calculated as the weight (kg) divided by the square of height (m). Patients with a BMI over or equal to 25 kg/m² were considered as overweight. Waist and hip circumferences were measured using a measuring tape accurate to 0.1 cm. Measurements were performed by a trained investigator during the morning under fasting conditions.

Body composition was estimated by bioelectrical impedance analysis (Tanita BC 420 S MA, Tanita Corp., Tokyo, Japan). Body fat percentage (%) and fat-free mass (kg) were assessed. Reference values of body fat percentage and fat-free mass in healthy European subjects [45] were used to estimate if individuals had excess body fat and a lack of fat-free mass according to their age and sex. Participants with excess body fat despite exhibiting a normal BMI (< 25 kg/m²) were considered as “normal weight obese” [46].

### Adherence to the French nutritional guidelines

The validated Programme National Nutrition Santé Guideline Score (PNNS-GS) was used to consider the adherence of individuals to the French dietary guidelines for fruits and vegetables, starchy foods, milk and dairy, meat, fats, sweetened foods, beverages, salt intake, and PA [47]. The maximum score was 15.

### Metabolic syndrome

If a participant possessed three or more of the following risk factors, they were considered to have metabolic syndrome according to the IDF and AHA/NHLBI thresholds [48]. Risk factors included: a waist circumference ≥ 94 cm in men and ≥ 80 cm in women; low HDL < 1.03 mmol/L in men and < 1.29 mmol/L in women; elevated triglycerides ≥ 1.7 mmol/L; elevated fasting glycemia ≥ 5.6 mmol/L; and elevated blood pressure (systolic blood pressure ≥ 130 mmHg and diastolic blood pressure ≥ 85 mmHg).

### Statistical analysis

The baseline characteristics of participants were described by gender and fertility status (frequency and percentage of categorical variables, as well as the mean and standard deviation of quantitative variables). Men and women were analysed separately due to their differences in lifestyle and body composition, as well as the different physiological benefits of exercise in both genders [49–51]. Comparisons between case and controls were conducted using Fisher’s exact test (as appropriate for categorical variables) and independent *t*-test (for continuous variables). Pearson correlation coefficients were computed to assess the relationship between PA level and sedentary behavior, and between PA level and body fat percentage. Analyses were performed separately for men and women. Associations between PA and sedentary behaviour with fertility status were investigated using logistic regression models. Multivariable analyses were performed after crude univariate logistic regressions. We elected not to include more than six covariates in the final model (age, education level, PA level, sedentary behavior, body fat and fat-free mass) in accordance with the literature [49] and to adhere to the principle of one variable studied for ten cases in small sample study. The regression model was adjusted for these six variables. Unadjusted and adjusted odds ratios (ORs) and 95% confidence intervals (CIs) were reported. BMI and waist circumference was not included in the models due to collinearity with body composition. SAS version 9.1 (SAS institute, Cary, NC, USA) was used to perform for all statistical analyses. A *p*<0.05 was considered significant.

## Results

Baseline characteristics of the 302 participants are presented in Table 1. Infertile participants were younger in comparison to fertile men and women (*p*=0.006 and *p*=0.02, respectively). They also had lower educational levels than fertile men and women (*p*=0.005 in men and *p*=0.0001 in women). The weight, BMI, waist circumference, hip circumference, and body fat of infertile men and women were significantly higher compared to fertile men and women. In men, the proportion of participants with metabolic syndrome was higher in infertile compared to fertile participants [12 (16.0%) *vs.* 3 (4.4%), respectively *p*=0.03]. The proportion of normal weight obese did not differ between groups of fertile and infertile men [5 (6.9%) and 5 (6.3%), respectively, *p*=1] and women [22 (31.0%) and 20 (25.0%), respectively, *p*=0.5)]. Mean PA levels did not significantly differ between fertile and infertile men (2726.2 and 3291.2 MET-min/week, respectively, *p*=0.2) and women (2632.8 and 2769.4 MET-min/week, respectively, *p*=0.3). However, infertile men spent more time performing moderate PA (121.0±181.2 min/week *vs.* 88.7±110.9 min/week, *p*<0.0001) and less time performing vigorous PA (37.6 ±48.6 min/week *vs.* 69.3±84.4 min/week, *p*<0.0001) in comparison to fertile men. Mean walking time (42.3±73.8 min/week *vs.* 35.9±36.4 min/week, *p*<0.0001) was higher in infertile men compared to fertile men, while it was lower (29.7±34.4 min/week *vs.* 46.6±66.5 min/week, *p*<0.0001) in infertile women compared to fertile women. Physical activity was only inversely associated with sedentary behavior in infertile men (r_Pearson_=-0.3, *p*=0.04). Physical activity was only inversely associated with body fat percentage in fertile men (r_Pearson_=-0.3, *p*=0.03). All infertile and fertile participants followed nutritional guidelines similarly, with scores of 6.6±2.1 vs. 6.2±1.9, respectively, for men (*p*=0.4), and scores of 6.3±2.9 vs. 6.3±3.1, respectively, for women (*p*=0.6) (maximal possible score of 15). Based on PA guidelines, 34 (47.2%) and 50 (63.3%) (*p*=0.05) fertile and infertile men did not follow PA guidelines (150 min/week of moderate to vigorous PA), respectively. Moreover, 43 (60.6%) and 55 (68.8%) (*p*=0.3) fertile and infertile women were under the recommended PA level, respectively.

**Table 1.**
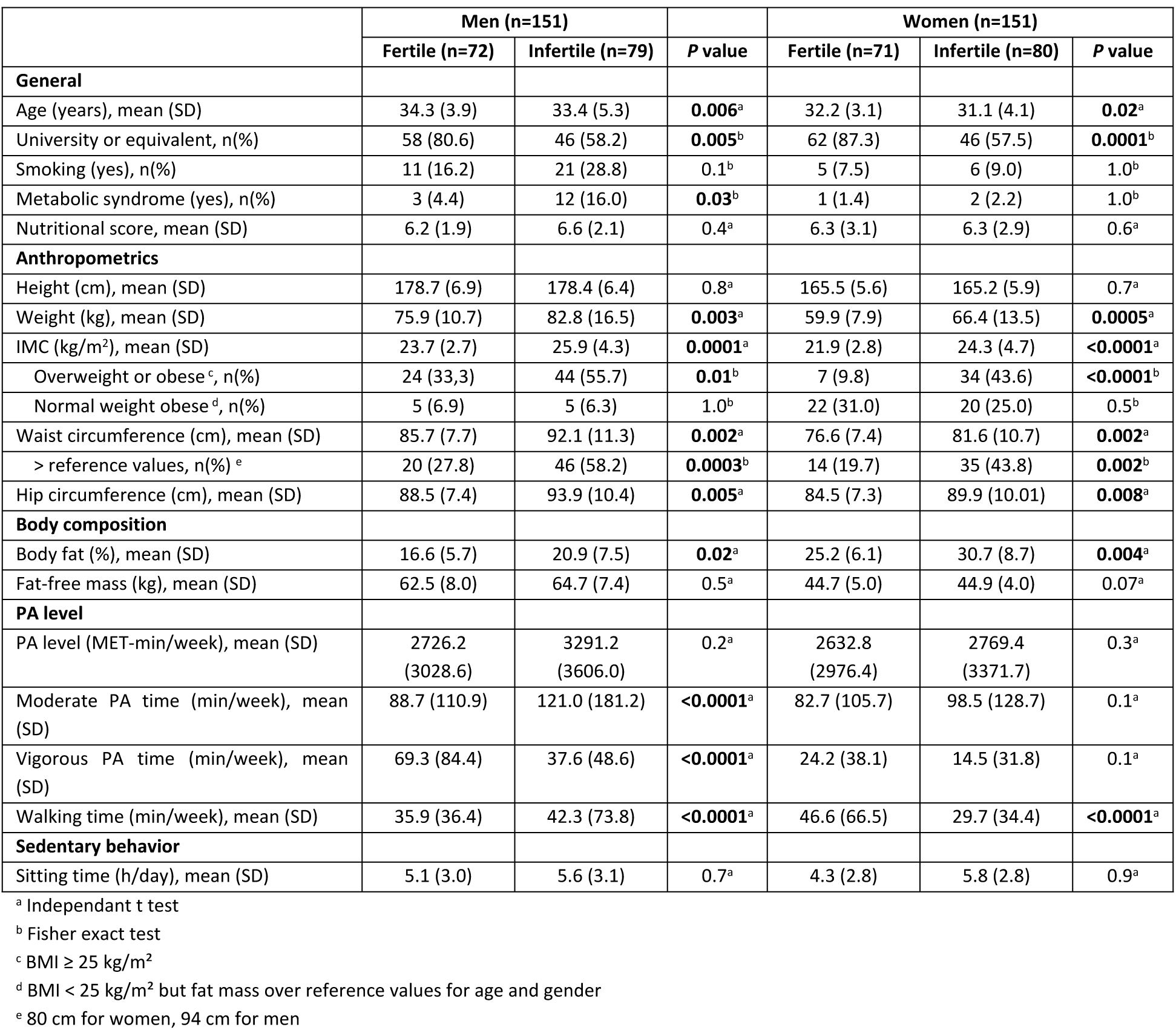
Baseline characteristics of fertile and infertile men and women

PA, sedentary behavior, and body composition factors according to fertility status and gender are presented in Table 2. In men, being physically inactive (adjusted OR 2.20; 95% CI, 1.06, 4.58; *p*=0.04) and having excess body fat (adjusted OR 2.83; 95% CI, 1.31, 6.10; *p*=0.008) were positively associated with infertility. Sedentary behavior and fat-free mass were not related to infertility in men in our study. In women, exhibiting sedentary behavior (adjusted OR 3.61; 95% CI, 1.58, 8.24; *p*=0.002) and possessing body fat over (adjusted OR 3.16; 95% CI, 1.36, 7.37; *p*=0.008) and fat-free mass under (adjusted OR 2.65; 95% CI, 1.10, 6.37; *p*=0.03) reference values for their age were associated with a significantly increased risk of infertility. Physical activity was not significantly associated with fertility status among women in our study (*p*=0.3).

**Table 2.** Factors associated with fertility and infertility (multivariable logistic regression)

|  | Men (n=151) |  |  |  | Women (n=151) |  |  |  |
| --- | --- | --- | --- | --- | --- | --- | --- | --- |
|  | % |  | Model |  | % |  | Model |  |
|  | Fertile<br>(n=72) | Infertile<br>(n=79) | OR [95% CI] | Adj OR<sup>a</sup> [95% CI] | Fertile<br>(n=71) | Infertile<br>(n=80) | OR [95% CI] | Adj OR<sup>a</sup> [95% CI] |
| **PA level (%)** |  |  |  |  |  |  |  |  |
| ≥150 min/week | 52.8 | 36.7 | 1.00 | 1.00 | 39.4 | 31.2 | 1.00 | 1.00 |
| <150 min/week | 47.2 | 63.3 | 1.93 (1.01-3.69) | 2.20 (1.06-4.58) | 60.6 | 68.8 | 1.43 (0.73-2.80) | 1.58 (0.73-3.42) |
| **Sedentary behavior (%)** |  |  |  |  |  |  |  |  |
| <5 h/day | 45.8 | 53.2 | 1.00 | 1.00 | 63.4 | 50.0 | 1.00 | 1.00 |
| ≥5 h/day | 54.2 | 46.8 | 0.75 (0.39-1.41) | 1.20 (0.55-2.61) | 36.6 | 50.0 | 1.73 (0.90-3.32) | 3.61 (1.58-8.24) |
| **Body fat (%)** |  |  |  |  |  |  |  |  |
| normal | 70.8 | 45.6 | 1.00 | 1.00 | 60.5 | 35.0 | 1.00 | 1.00 |
| > ref. values<sup>b</sup> | 29.2 | 54.4 | 2.90 (1.48-5.69) | 2.83 (1.31-6.10) | 39.4 | 65.0 | 2.85 (1.47-5.53) | 3.16 (1.36-7.37) |
| **Fat-free mass (%)** |  |  |  |  |  |  |  |  |
| normal | 55.6 | 65.8 | 1.00 | 1.00 | 66.2 | 62.5 | 1.00 | 1.00 |
| < ref. values<sup>b</sup> | 44.4 | 34.2 | 0.65 (0.34-1.25) | 0.89 (0.42-1.87) | 33.8 | 37.5 | 1.75 (0.60-2.29) | 2.65 (1.10-6.37) |

Abbreviations: OR, Odds ratio, Adj OR, Adjusted Odds ratio, CI, Confidence Interval, PA, Physical Activity, SD, Standard Deviation.
<sup>a</sup> Adjusted for age and educational level and for all variables of the table.
<sup>b</sup> Age and gender reference values [45].

## Discussion

Idiopathic infertility in men and women may be related to lifestyle and body composition factors. In this case-control study, physical inactivity in men and sedentary behavior in women were independently associated with infertility. Body fat accumulation was significantly independently associated with infertility status in both men and women, while fat-free mass was related to infertility only in women.

Consistent with our study, sedentary behavior has not been significantly associated with infertility or related factors in men [18,24,25]. However, higher volumes of television viewing time have been related to vitamin D deficiency [52], which has been associated with a lower percentage of motile spermatozoa compared to men with sufficient vitamin D levels [32,53]. Low levels of vitamin D are also related to obesity [54,55]. Adiposity, often associated with sedentary behavior and physical inactivity [56], increases oxidative stress, which may subsequently lead to gonad and gamete damage in men [57]. Regarding the men in our study, being over the reference values for body fat was independently related to infertility. We also note that metabolic syndrome was more frequent among infertile men than fertile men in our study. Metabolic syndrome—as well as oxidative stress related to obesity—have been associated with reductions in sperm concentration, count, motility, and vitality [58,59].

Physical inactivity was related to infertility in men, independently from sedentary behavior. We have also demonstrated that less PA was resulted in more sedentary behavior being present in infertile men. Leisure PA, specifically outdoor and weight lifting activities, have been associated with higher sperm concentration in a dose-response relationship, though was not associated with higher reproductive success in the context of fertility treatment [10]. It has been observed that men who are moderately active three times per week for one hour had better sperm morphology in comparison to men who participated in more intense and frequent PA or cycling activities [7]. However, our population is not comparable to Vaamonde *et al.*’s study, as they spend a mean time of less than 2 hours per week engaging in moderate PA. Moreover, while intense leisure PA has been associated with lower sperm quality [18], fertile men in our study spent more time engaging in vigorous PA than infertile men. As a recent study highlighted, different types of PA may affect semen quality parameters differently [9]. As such, further studies are needed to investigate the volume of PA that may be specifically related to male fertility.

Among women, sedentary behavior was associated with infertility, while other studies were unable to confirm a significant relationship between this behavior and fertility as well as probability of live birth [17,26]. Sedentary behavior has been positively associated with the secretion of leptin [60], which can decrease fertility [56] and pregnancy rates with *in vitro* fertilization (IVF) through the downregulation of the hypothalamic-pituitary-ovarian (HPO) axis [61]. In turn, this downregulation of HPO affects gonadotropin production, which may lead to menstrual abnormalities and ovulation dysfunction [30].

Body fat was independently associated with fertility among women in our study, and this seems to be a confounding factor of sedentary behavior effects on proinflammatory cytokine regulation [56,61,62]. Notably, sedentary behavior is independently associated with central adiposity [63] and total adiposity [56]. This deleterious accumulation of fat is important in the production of adipocytokines, which influence estrogen biosynthesis [64]. Adiposity may also compromise the reproductive endocrine system through increased androgen and estrogen secretion, as well as decreased sex hormone binding globulin (SHBG) secretion [56,62]. The link between body fat and fertility was also described in infertile women with ovarian failure [42]. We did not test the association of fertility with overweight status through BMI since this relationship is well established, and because BMI is a somewhat crude measure of body fatness; instead, we had access to actual body composition measures [29]. Moreover, normal weight obesity should be considered. If we focus on the proportion of infertile women with high body fat and high BMI in addition to the proportion of normal weight obese women (with high body fat but a normal BMI), we obtained a total of 68.6%. This high proportion of infertile women at risk due to their body composition should be considered in future research.

Notably, a low fat-free mass was positively associated with infertility in women. To our knowledge, this relationship has been poorly documented. Kirchengast *et al.* have highlighted a positive relationship between infertility and a reduced bone mineral content, but not with lean body mass [42]. Another study found no statistical association between infertility and bone mineral density and lean body mass [65]. Therefore, we can suggest that lean body mass could be implicated in infertility since it plays an important role in the control of systemic energy metabolism and insulin sensitivity [66]—both of which interfere with fertility [67]. Moreover, low muscle mass can be associated with oxidative stress [68]. Bone mineral content has also been associated with decreased sex hormone levels [69].

In line with other studies, we did not confirm a significant association between physical inactivity and fertility in women [17,26]. However, it appears that the relationship between PA and fertility may differ according to BMI [17]. In opposition to our results, Wise *et al*. demonstrated that in women, moderate PA increased fecundity parameters independently of BMI [13]. Gudmundsdottir *et al.* also described a u-shaped relationship between duration of exercise and infertility in younger women (less than 30 years old). Subgroups of women exercising under 15 min and over 60 min per session had a higher frequency of infertility than women between 16 and 60 min duration [21]. PA was not related to fertility status among women in our study, likely due to a lack of power; however, it has been shown that 1.5 h or more of aerobic PA per week resulted in a higher likelihood of live birth in women during IVF compared to inactive women [17]. As it has been observed in men, total PA level (in MET-min/week) may not be the variable most highly associated with fertility status, while PA parameters such as duration, intensity, frequency, and type of exercise could be instead. This decomposition, known as the FITT principle (i.e. frequency, intensity, time, and type of exercise) [70], should be studied further in association with fertility in men and women.

While useful, this study presents some limitations. Our findings have to be nuanced while the study may present lack of power to detect associations. Furthermore, our findings are limited to men and women with unexplained infertility. Consequently, we were unable to observe the consequences of a lack of PA or high sedentary behavior on conventional sperm parameters or ovarian failure. Physical activity was estimated through a self-assessment questionnaire. The risk of declarative data on PA and sedentary behavior on the last-7-day recall must be taken in consideration because it can increase the risk of over-or under-estimation [71]. PA level corresponds to usual PA at the time of inclusion, though data on lifetime PA and variations in PA over time should be studied further. Notably, PA history during adulthood has been positively associated with PA level [49]. Thus, we can expect that, in our population, current PA behaviors are reflective of PA behavior in the recent past. Moreover, the IPAQ questionnaire does not allow for the investigation of FITT parameters, which would have provided more specific data on associations between the specific dimensions of PA and infertility. Despite these limitations, the IPAQ questionnaire has been validated in various populations against accelerometer and pedometer data [71]. Bioelectrical impedance analysis is not the gold standard measurement of body composition, nor is it as specific as dual-energy X-ray absorptiometry. In the regression model, we elected to use dichotomous variables in order to be over or under the recommended PA level, 5h of sedentary behavior, and reference values for body fat and fat-free mass in order to have parameters that are easily operational in clinical practice; however, the actual threshold may involve less discriminant criteria. In a recent study, PA and sedentary behavior analyzed on a continuous level did not show associations with fertility [17]. In our study, post-hoc analyses have also demonstrated no association (data not shown). Infertile couples were selected by medical services, while the control group consisted of fertile volunteers recruited from the general healthy population within the areas of participating medical services. Case and control groups were comparable regarding most variables other than the study variables, and the assessments were performed for all participants using the same trained investigator. Despite efforts to recruit comparable subjects, we observed differences between case and control groups in terms of socio-economic status. Our model has been adjusted for educational level, which may limit the impact of this difference. Fertile participants were slightly older than those that were infertile, which could be explained by the fact that they were recruited after the birth of their child and they were not usually included immediately after childbirth. The study of couple-based associations was not possible and it should be explored further in future studies.

These findings of the present study suggest that sedentary behavior and physical inactivity would be two independent factors to consider regarding fertility, as has been suggested for the general population [4]. Beyond the type of exercise performed, it appears that the frequency, duration, intensity and the type of PA may affect infertility parameters differently in men and women [10,17]. Further investigations on the FITT criteria of PA should be undertaken in order to propose recommendations. Moreover, sedentary behavior should be more widely investigated. In particular, sedentary behavior must be studied in relation to its accumulation process, since interrupting the amount of time spent sitting was related to decreased visceral adiposity and BMI [72,73]. Further studies on the interaction of sedentary and PA behaviors regarding fertility is also warranted. Additionally, the relationship between fat and fat-free mass with fertility may be of interest. Studies have recently been implemented in obese rats to explore the relationship between body composition and reproductive programming through oxidative stress regulation while training [74,75]. It has been suggested that the amount and distribution of fat and lean tissue may influence reproductive factors differently [76]. The relationship between semen parameters among infertile men of the ALIFERT study and PA, sedentary behavior and body composition is currently under investigation. In addition to the usual care for infertility treatment, an improvement in major modifiable lifestyle factors should be considered. A prospective interventional randomized controlled trial would be relevant to test this hypothesis. Meanwhile, practical advice and education might already be proposed, such as regularly being physically active and breaking up sedentary behavior time [6,77].

## Conclusion

The present study demonstrated that physical inactivity in men and sedentary behavior in women are associated with idiopathic infertility. Body fat accumulation has been related to infertility in both men and women, while fat-free mass was related to infertility in women only. This case-controlled study highlights that physical inactivity and sedentary behavior represent two independent risk factors for infertility. The effect of various elements that make up PA (i.e. FITT criteria) and interrupting the time spent sitting were not tested in this study, and should be considered in future research. The differences observed between men and women should also be studied further. Moreover, body composition variation through lifestyle should be also explored further in relation to the biological pathways involved in idiopathic infertility. These findings suggest promoting and proposing a lifestyle supportive care during fertility treatment in order to improve pregnancy rates.

## Author’s roles

AMF, CD, RL and CJ participated in analysis, interpretation and manuscript drafting. AMF and CJ performed statistical analysis. CD, CF and the ALIFERT collaborative group participated in the recruitment of patients. RL, CD, SC, CF and the ALIFERT collaborative group participated in the conception, the design, and the execution of the study. AMF, CF, CJ, SC, RL, and CD participated in critical discussion of the study for intellectual content and gave a final approval of the manuscript.

## Acknowledgements

The authors acknowledge all the couples involved in the study and the ALIFERT collaborative group.

## Supporting information

**S1 Table. Baseline characteristics of fertile and infertile men and women.** ^a^ Independant t test, ^b^ Fisher exact test, ^c^ BMI ≥ 25 kg/m², ^d^ BMI < 25 kg/m² but fat mass over reference values for age and gender, ^e^ 80 cm for women, 94 cm for men.

**S2 Table. Factors associated with fertility and infertility (multivariable logistic regression).** Abbreviations: OR, Odds ratio, Adj OR, Adjusted Odds ratio, CI, Confidence Interval, PA, Physical Activity, SD, Standard Deviation, ^a^ Adjusted for age and educational level and for all variables of the table, ^b^ Age and gender reference values [45].

